# Labile iron pool dynamics do not drive ferroptosis potentiation in colorectal cancer cells

**DOI:** 10.1101/2025.07.01.662602

**Authors:** Varun Ponnusamy, Deahzana R. Randall, Zheng Hong Lee, Nupur K. Das, Liang Zhao, Kathryn Buscher, Sumeet Solanki, Adam R. Renslo, Peggy P. Hsu, Yatrik M. Shah

**Affiliations:** Department of Molecular and Integrative Physiology, University of Michigan, Ann Arbor, MI, USA.; Graduate Program of Immunology, University of Michigan, Ann Arbor, MI, USA; Department of Pharmaceutical Chemistry, University of California, San Francisco, San Francisco, California, USA; Rogel Cancer Center, University of Michigan Medical School, Ann Arbor, MI, USA; Department of Internal Medicine, Division of Hematology and Oncology, University of Michigan Medical School, Ann Arbor, MI, USA; Department of Internal Medicine, Division of Gastroenterology, University of Michigan Medical School, Ann Arbor, MI, USA

**Keywords:** Iron, Metal Homeostasis, Cell Death, Oxidative Stress, Lipid Peroxidation, Colon Cancer, Metabolic Regulation

## Abstract

Colorectal cancer (CRC) is the second leading cause of cancer-related mortality in the United States. CRC tumors exhibit aberrant iron accumulation, which supports tumor cell proliferation through multiple metabolic pathways. However, the oncogenic benefits of elevated iron must be counterbalanced by its potential to catalyze oxidative damage via reactive oxygen species generated from labile, redox-active iron. Ferroptosis is a regulated, non-apoptotic form of cell death characterized by iron-dependent lipid peroxidation. This process is tightly controlled by the selenoenzyme glutathione peroxidase 4 (GPX4), which reduces lipid peroxides and can be pharmacologically inhibited by agents such as RSL3 and JKE1674. A key source of redox-active iron is the labile iron pool (LIP), yet its role in regulating ferroptosis remains incompletely defined. To examine this, we supplemented CRC cells with exogenous iron following pharmacologic induction of ferroptosis. Iron supplementation significantly reduced cell viability, suggesting that expansion of the LIP potentiates ferroptotic cell death. However, whether ferroptosis is accompanied by dynamic changes in the LIP, and if such changes are mechanistically required for its potentiation, were unknown. To further characterize this response, we profiled the expression of iron regulatory genes under ferroptotic conditions and observed no change in transcriptional response in iron homeostasis genes. Using a reactivity-based probe of labile iron, we found that the LIP did not measurably increase during ferroptosis induction with GPX4 inhibition or inhibition of the SLC7A11 cysteine/glutamate antiporter. These findings suggest that the LIP does not expand upon pharmacological initiated ferroptosis, despite the potentiating effect of exogenous iron supplementation.

## Introduction

Iron is an essential micronutrient that supports a wide range of cellular processes^1^. Given that both iron deficiency and overload can be harmful, iron levels are tightly regulated. The intestine plays a central role in this regulation, as it is the sole site of dietary iron absorption and the gateway for systemic iron distribution to peripheral tissues. Intestinal epithelial cells are exposed to two principal sources of iron: luminal dietary iron and iron derived from the systemic circulation. In the intestine, iron enters epithelial cells apically via divalent metal transporter 1 (DMT1; Slc11a2) after reduction to the ferrous (Fe^2+^) state, or basolaterally in its ferric (Fe^3+^) transferrin-bound form through the transferrin receptor (TFRC)^2,3^. Once inside the cell, iron is reduced to its ferrous form and is either incorporated into iron-dependent enzymes and cofactors or trafficked to the mitochondria for utilization^4^. Due to its redox reactivity and potential to generate reactive oxygen species (ROS), intracellular iron is tightly controlled. Excess iron is either exported by the basolateral iron transporter ferroportin (FPN)^5^ or stored within the protein complex ferritin (FTN)^6^. Under conditions of increased demand, stored iron can be mobilized through ferritinophagy, a selective autophagic process mediated by the cargo receptor nuclear receptor coactivator 4 (NCOA4)^7,8^. A central component of cellular iron regulation is the labile iron pool (LIP). The LIP is a cytosolic, redox-active, and readily chelatable pool of ferrous iron that serves as the key bioavailable iron reservoir.

A form of regulated, non-apoptotic cell death, known as ferroptosis, is characterized by its dependence on iron and its initiation through the accumulation of lipid peroxides^9,10^. The LIP plays a central role in ferroptosis by catalyzing the decomposition of hydrogen peroxide and oxygen via the Fenton reaction^11^, generating highly reactive hydroxyl and hydroperoxyl radicals. These ROS cause oxidative damage to cellular membranes, particularly through the peroxidation of polyunsaturated fatty acids in membrane phospholipids, ultimately compromising membrane integrity and triggering ferroptotic cell death^12^.

Ferroptosis research thus far has relied on synthetic small molecules known to induce ferroptotic cell death. Only recently have endogenous regulators of ferroptosis begun to be identified^13^. There remains a need to better understand endogenous mechanisms that mediate ferroptosis, including the role of iron itself, an underexplored but potentially critical regulator. Notably, certain cancers, such as colorectal cancer (CRC), exhibit a strong dependence on iron for growth, and are potentially susceptible to ferroptosis-based therapies^14–17^. Early work suggested that LIP is increased during ferroptosis; however, these studies^18,19^ employed chelation-based iron sensors that lack intrinsic metal ion and oxidation-state specificity. Gaining deeper insight into ferroptosis and the role of iron in its regulation may offer novel therapeutic strategies for CRC and other iron-dependent cancers.

In this study, we demonstrate that supplementation with exogenous free iron significantly enhances ferroptosis when combined with established ferroptotic inducers in multiple human CRC cell lines. To determine whether a concordant increase in LIP potentiates ferroptosis via endogenous mechanisms, we investigated whether the LIP is naturally altered during ferroptosis. This was accomplished by profiling the expression of key iron homeostasis genes and using TRX-PURO, a reactivity-based probe with high sensitivity and Fe^2+^ specificity, to detect subtle changes in the LIP. Interestingly, our analysis revealed that, at the detection limits of TRX-PURO, no significant increase in endogenous LIP occurs during ferroptosis. These findings suggest that, contrary to expectations, the LIP remains relatively stable in the absence of exogenous iron manipulation and that additional regulatory mechanisms beyond iron availability may be required to effectively drive ferroptotic cell death.

## Results

### Exogenous iron sensitizes CRC cells to JKE-induced ferroptosis

The selenoprotein glutathione peroxidase 4 (GPX4) is a central regulator of ferroptosis^20^. Although inhibition of GPX4 is widely accepted as a mechanism to induce ferroptosis, we have previously shown that the commonly used GPX4 inhibitor RSL3 exhibits off-target effects and non-specific inhibition across the selenoproteome, including thioredoxin peroxidases^21^. We therefore utilized a more specific, next-generation GPX4 inhibitor, JKE1674, for our experiments.

Among the CRC cell lines, HCT116 and SW480 have been reported to be relatively resistant to ferroptosis inducers, whereas RKO is highly sensitive^17^. We found that JKE1674-mediated GPX4 inhibition was less potent in HCT116, SW480, and RKO compared to other ferroptosis inducers^22^ (Figure 1A, S1A). To determine whether manipulating the labile iron pool could sensitize cells to JKE1674 (0.625, 1.25, 2.5 μM), we co-treated cells with ferric ammonium citrate (FAC) at increasing concentrations (250, 500, and 1000 μM). The fibrosarcoma cell line HT1080, a well-established ferroptosis sensitive cell line^13^, was included as a positive control. Exogenous iron significantly reduced cell viability in the ferroptosis-resistant CRC lines HCT116 and SW480, particularly at the highest FAC concentration (Figure 1B).

**Figure 1:**
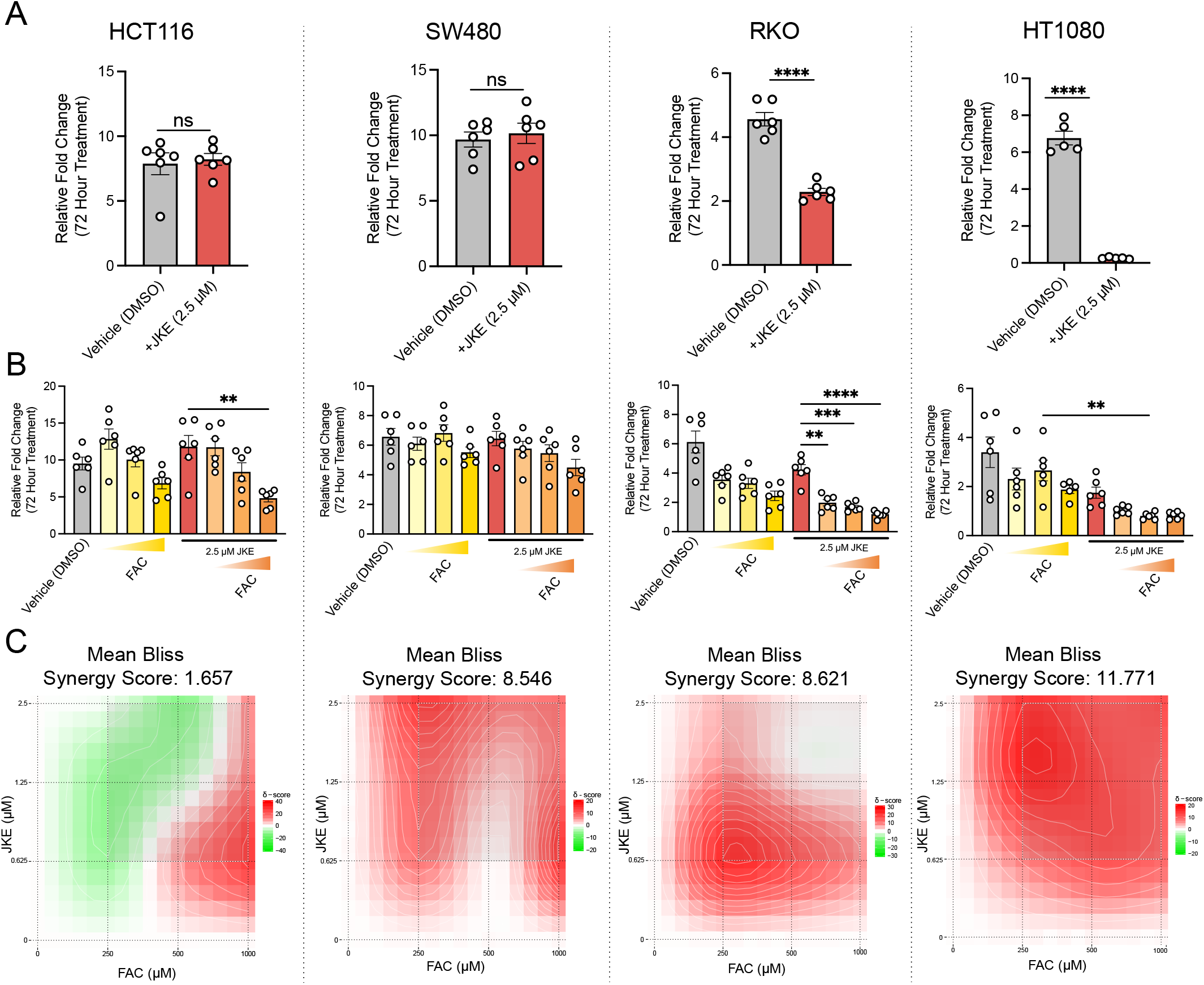
Exogenous iron sensitizes CRC cells to JKE-induced ferroptosis. (A) Cell viability of HCT116, SW480, RKO, and HT1080 cells following treatment with 2.5 μM JKE1674 after 72 hours (B) Cell viability of HCT116, SW480, RKO, and HT1080 cells following treatment with 2.5 μM JKE1674 with the addition of 250, 500, 100 μM FAC after 72 hours (C) Heatmap plot of synergy in HCT116, SW480, RKO, and HT1080 cells treated with JKE1674 and FAC ∗p < 0.05, ∗∗p < 0.01, ∗∗∗p < 0.001, ∗∗∗∗p < 0.0001 unless otherwise indicated. Data are presented as mean ± SEM. p < 0.0001 unless otherwise indicated. Data are presented as mean ± SEM.

To quantify the interaction between FAC and JKE1674, we performed synergy analysis using the Bliss independence dose-response surface model^23^. This model assumes additivity when two agents act independently, with deviations reflecting synergy or antagonism. Synergy scores >10 indicate synergy, scores between –10 and 10 indicate additivity, and scores <–10 indicate antagonism. The mean synergy score represents the average of all values depicted in the heatmap. Across all four cell lines tested, growth assays revealed robust synergistic interactions between FAC and JKE1674 over a range of concentrations (Figure 1C).

### Exogenous iron potentiates RSL3 and IKE-induced ferroptosis in CRC cells

Next, we tested whether FAC could potentiate two other ferroptotic inducers, RSL3 and the SLC7A11 inhibitor, imadazole ketone erastin (IKE) (Figure 2A, 2B). Co-treatment of HCT116 and RKO cells with RSL3 (1, 0.5, 0.25 μM) and FAC resulted in a dose-dependent decrease in cell viability. In contrast, SW480 cells, which are highly resistant to ferroptosis, did not exhibit enhanced sensitivity to either RSL3 or IKE upon FAC addition.

**Figure 2:**
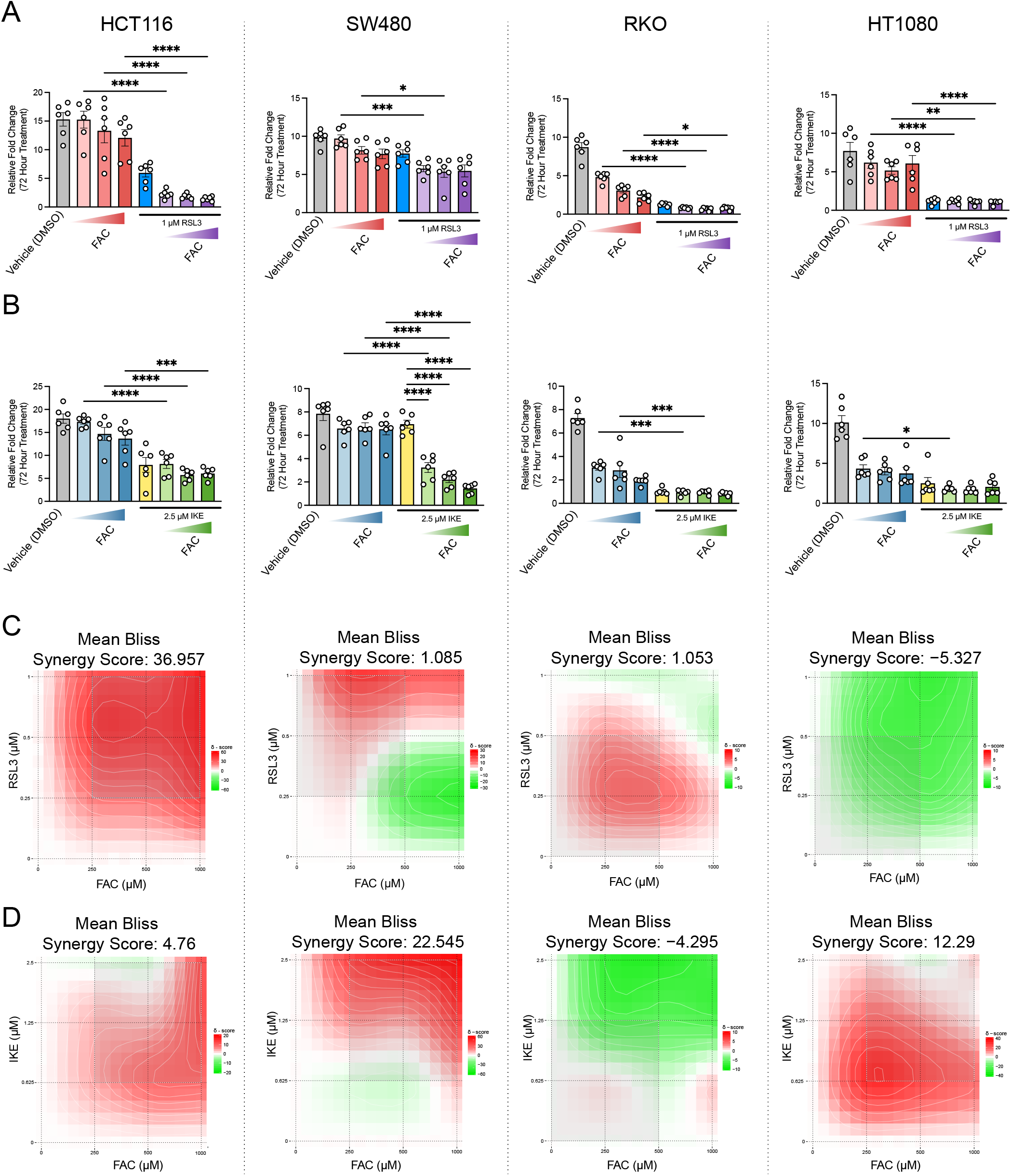
Exogenous iron potentiates RSL3 and IKE-induced ferroptosis in CRC cells. (A) Cell viability of HCT116, SW480, RKO, and HT1080 cells following treatment with 1 μM RSL3 after 72 hours (B) Cell viability of HCT116, SW480, RKO, and HT1080 cells following treatment with 2.5 μM IKE with the addition of 250, 500, 1000 μM FAC after 72 hours (C) Heatmap plot of synergy in HCT116, SW480, RKO, and HT1080 cells treated with RSL3 and FAC (D) Heatmap plot of synergy in HCT116, SW480, RKO, and HT1080 cells treated with IKE and FAC ∗p < 0.05, ∗∗p < 0.01, ∗∗∗p < 0.001, ∗∗∗∗p < 0.0001 unless otherwise indicated. Data are presented as mean ± SEM.

Ferroptosis can also be triggered by inhibition of the cystine/glutamate antiporter SLC7A11, which increases lipid peroxidation^9,24^. Notably, FAC potentiated IKE-induced cell death in the SW480 cell line. HCT116 cells showed a trend toward decreased viability at lower IKE concentrations, similar to effects seen in RKO and HT1080 cells (Figure 2B, S2B).

To quantify these interactions, we again applied the Bliss independence dose-response surface model. Co-treatment with RSL3 and FAC demonstrated strong synergy in HCT116 cells and additive effects in the remaining lines (Figure 2C). For IKE, synergistic effects were observed in SW480 and RKO, with additive responses in HCT116 and HT1080 (Figure 2D). Together, these results show that increasing the labile iron pool through exogenous iron supplementation can potentiate ferroptosis both through GPX4-dependent and GPX4-independent mechanisms.

### Iron homeostatic genes are not altered following RSL3, JKE, and IKE-induced ferroptosis

In the intestine, iron is imported and exported through the key transporters DMT1, TFRC, and FPN (the only known mammalian iron exporter). FTN serves as the primary intracellular iron storage protein, sequestering excess iron from the LIP^1,25^. The expression of these iron homeostasis genes is regulated by iron regulatory proteins (IRPs), a class of RNA-binding proteins that control translation by binding to iron-responsive elements (IREs) within the untranslated regions (UTRs) of target mRNAs^26,27^. Depending on the location of IREs in the 3′ or 5′ UTR, IRP binding can upregulate or downregulate the expression of DMT1, TFRC, FPN, and ferritin heavy chain (FTH1) during systemic iron deficiency or overload^28,29^.

In addition to IRPs, iron homeostasis genes are transcriptionally regulated by transcription factors. Hypoxia-inducible factor (HIF)-2α and nuclear factor erythroid 2–related factor (NRF)-2 bind to hypoxia response elements and antioxidant response elements, respectively, particularly under conditions of hypoxia or oxidative stress^14,30–33^. While these regulatory systems maintain iron balance under physiological conditions, it is unclear whether they become dysregulated and contribute to ferroptosis.

It is widely assumed, but not rigorously tested, that ferroptosis is accompanied by an endogenous increase in the LIP, which would sensitize cells to iron-dependent lipid peroxidation ^18,19^. To investigate this, we examined the expression of iron homeostasis genes (DMT1, TFRC, FTH1, and FPN) following induction of ferroptosis. Quantitative RT-PCR was performed 16 hours after treatment with three distinct ferroptosis inducers (Figure 3). Surprisingly, ferroptosis induction did not result in significant transcriptional changes in iron homeostasis genes, except in SW480 cells treated with RSL3 (Figure 3A) and in HT1080 cells treated with IKE (Figure 3B). Though these changes in gene expression are not consistent across dosages. These findings suggest that, despite the central role of iron in ferroptosis, the LIP may not undergo an endogenous increase during ferroptosis, and that transcriptional regulation of iron genes may not be a primary driver of ferroptosis sensitivity.

**Figure 3:**
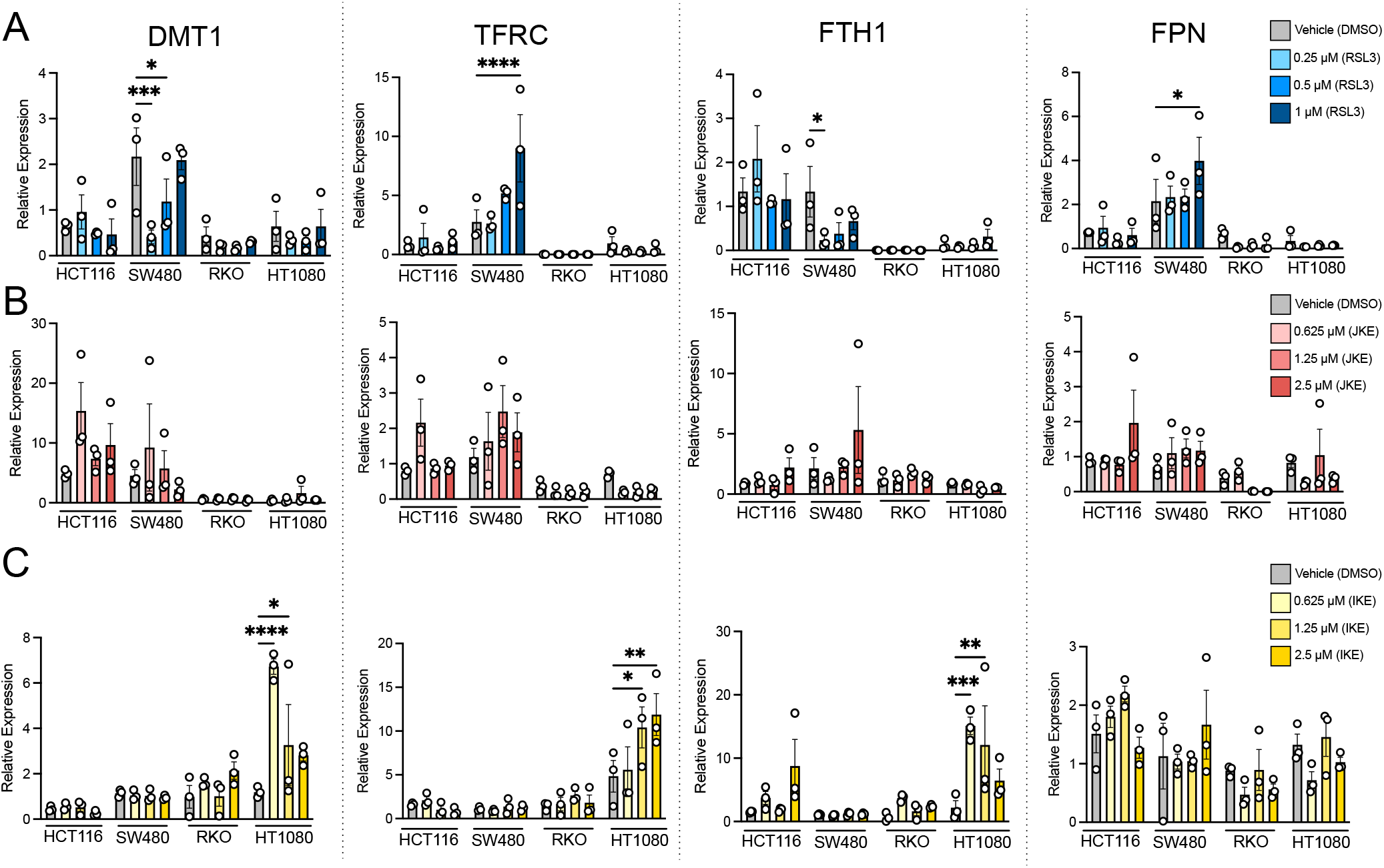
Iron homeostatic genes are not altered in RSL3, JKE, and IKE-induced ferroptosis in CRC cells. (A) qPCR of iron homeostasis genes in HCT116, SW480, RKO, and HT1080s treated with 0.25, 0.5, and 1 μM RSL3 after 16 hours (B) qPCR of iron homeostasis genes in HCT116, SW480, RKO, and HT1080s treated with 0.25, 0.5, and 1 μM JKE1674 after 16 hours

### The labile iron pool is not altered during ferroptosis

Early studies suggested that the LIP increases following the initiation of ferroptosis, primarily through the degradation of ferritin via ferritinophagy^18,19^. These studies measured the LIP using Calcein AM^34–36^ and PhenGreen SK^37^, chelation-based iron probes that, while still widely used to measure LIP, have inherent limitations, as has been reviewed^38^. Specifically, these probes are dependent on esterases to cleave lipophilic ester functions that promote cellular permeability, leading to variable cellular uptake and distribution. Moreover, since binding of analyte to these probes leads to fluorescence quenching, a second, cell-permeable iron chelator (typically 2,2’-bipyridine or 1,10-phenantrholine) is employed to produce a turn-on response through competition for chelation. Accordingly, the measured response of such probes is defined operationally, a function of the amount of probe loaded on cells, the extent of ester hydrolysis and intracellular sequestration, and the relative affinities of the probe and second chelator for Fe^2+^ and Fe^3+^. While quantitative measurement of LIP with such methods is theoretically possible, this requires careful *in situ* or *ex situ* calibration experiments that are rarely performed in practice.^36,39^

To assess LIP levels after ferroptosis induction with RSL3, JKE, or IKE, we employed the reactivity-based probe TRX-PURO,^39^ which detects LIP via the Fenton-type reactivity that is associated with labile Fe^2+^ but not with Fe^3+^ in storage nor with protein-bound iron co-factors. TRX-PURO is an endoperoxide-caged conjugate of puromycin that is uncaged upon reaction of the endoperoxide function with labile iron. Uncaged puromycin is incorporated into nascent polypeptides at the ribosome, where it can be readily detected by flow cytometry, western blotting, or immunofluorescence staining using α-puromycin antibodies (Figure 4A). We validated the iron-dependent response of TRX-PURO in HEK293 cells co-treated with the ferric iron source FAC (10–50 µM). FAC is only competent for TRX-PURO activation following its internalization and reduction to the ferrous LIP. Puromycin incorporation increased in a dose-dependent manner with FAC, consistent with enhanced labile iron availability (Figure 4B). Co-treatment with the iron chelator deferoxamine (DFO) reduced puromycin incorporation to baseline levels, confirming the probe’s specificity for iron.

**Figure 4:**
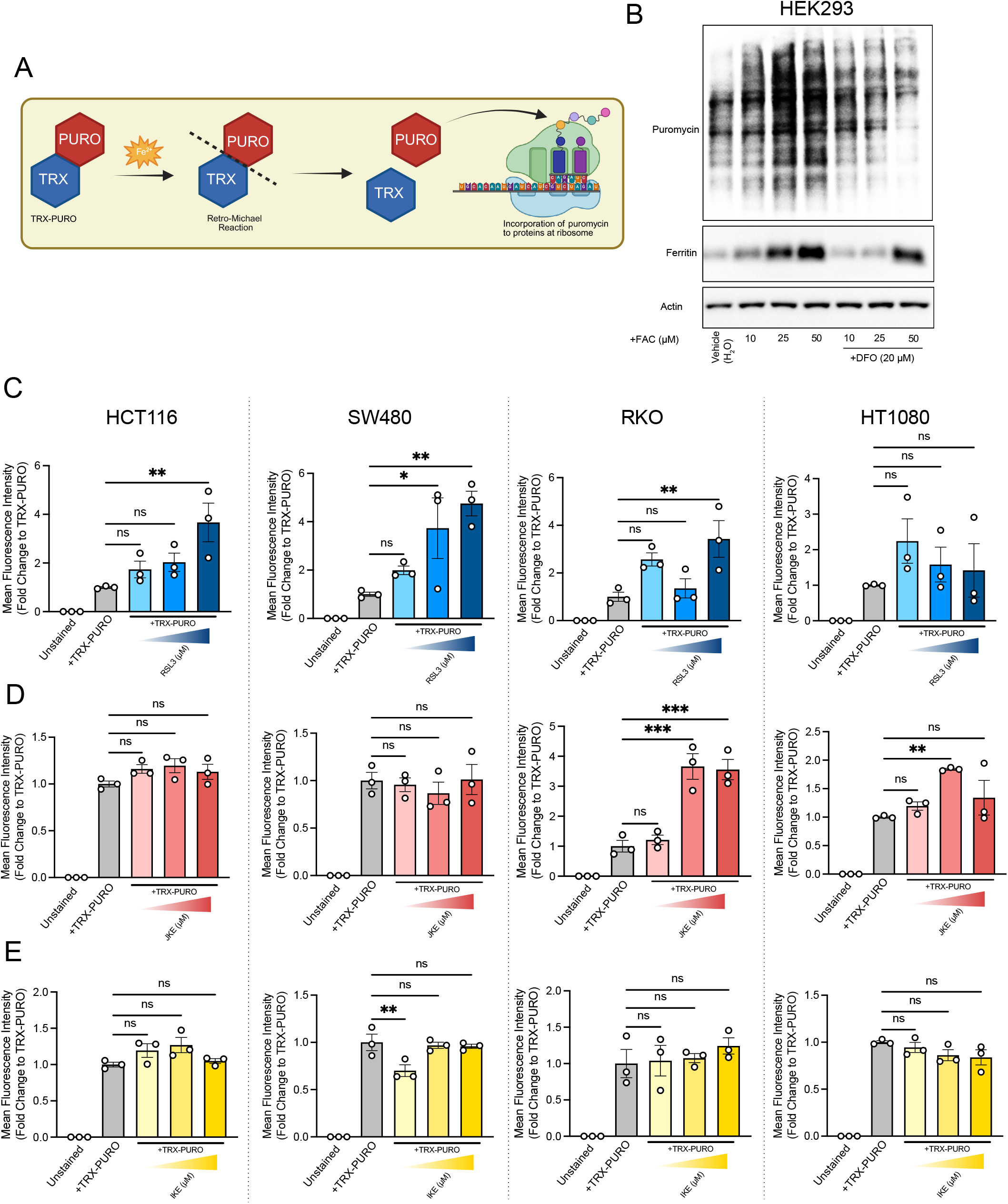
The labile iron pool is not altered during ferroptosis. (A) Schematic of mechanism of puromycin (PURO) release from TRX-PURO and incorporation into nascent proteins at ribosome (B) Validation of TRX-PURO in HEK293 cells co-treated with FAC and TRX-PURO by western blot to assess labile iron pool concentration (C) Quantification of fluorescent intensity of HCT116, SW480, HT1080, RKO cells stained with TRX-PURO (1 μM) to assess labile iron pool concentration after treatment of 1, 0.5, 0.25 μM RSL3 (D) Quantification of fluorescent intensity of HCT116, SW480, HT1080, RKO cells stained with TRX-PURO (1 μM) to assess labile iron pool concentration after treatment of 2.5, 1.25, 0.625 μM JKE (A) Quantification of fluorescent intensity of HCT116, SW480, HT1080, RKO cells stained with TRX-PURO (1 μM) to assess labile iron pool concentration after treatment of 2.5, 1.25, 0.625 μM IKE *p < 0.05, ∗∗p < 0.01, ∗∗∗p < 0.001, ∗∗∗∗p < 0.0001 unless otherwise indicated. Data are presented as mean ± SEM.

To determine whether the LIP increases endogenously during ferroptosis, we assessed TRX-PURO activation via puromycin incorporation over 16 hours of continuous treatment with probe in the presence of RSL3, JKE, or IKE (Figure 4C–E). Maximal activation of TRX-PURO requires ∼8 hours due to a rate-determining retro-Michael reaction involved in puromycin release^39^ and so these studies are necessarily limited in their ability to report on fluctuations in LIP at very early timepoints. Nevertheless, we found that increasing concentrations of RSL3 led to increased LIP in HCT116 and SW480 cells while RKO and HT1080 cells showed a more stochastic TRX-PURO that was not dose-responsive (Figure 4C and S3A). Conversely, JKE treatment did not affect LIP in HCT116 nor SW480 cells at any of the doses studied, while RKO and HT1080 cells showed elevated LIP at the higher doses (Figure 4D and S3B). Finally, IKE failed to increase labile iron over untreated controls in any of the cell lines examined (Figure 4E and S3C). Taken together, these data indicate that elevation of the LIP is not a consistently conserved or essential feature of pharmacologically induced ferroptosis.

## Discussion

Ferroptosis is a form of regulated cell death that depends on iron and is characterized by the accumulation of lipid peroxides^10^. Unlike apoptosis or necroptosis, ferroptosis lacks well-defined molecular markers. Additionally, whether endogenous mediators regulate ferroptosis is poorly understood. Among these regulators is thought to be iron and particularly the LIP. The LIP plays a central role by triggering the Fenton reaction and initiating lipid peroxidation. However, the extent to which endogenous fluctuations in iron availability through the LIP directly contribute to ferroptosis remains unknown.

Our data confirms that iron remains functionally essential for ferroptosis. When we increased intracellular iron levels through exogenous supplementation with FAC, CRC cells became markedly more sensitive to ferroptosis inducers. This synergy was observed in multiple ferroptosis-sensitive and -resistant CRC cell lines and the ferroptosis-sensitive HT1080 fibrosarcoma cell line.

Despite the reliance on iron for triggering ferroptosis our results suggest CRC cells may not potentiate ferroptosis through iron mobilization but instead exhibit a relatively stable LIP. Our quantification of the LIP using TRX-PURO indicates that higher concentrations of RSL3 endogenously increase the LIP in CRC cells. We have previously reported^21^ that RSL3 has several off targets independent of GPX4 which may contribute to an increase in LIP, making these results somewhat unreliable. When using JKE, a more specific GPX4 inhibitor, we saw no change in labile iron in ferroptosis-resistant cell lines, HCT116 and SW480 and increase in the LIP in ferroptosis-sensitive cell lines, RKO and HT1080 at higher JKE concentrations. IKE-induced ferroptosis showed no change to the LIP in any of our tested cell lines. This is in contrast to a study that showed erastin-induced ferroptosis increases the LIP in breast cancer cells^42^. The variation among cell lines implies context-specific regulation of iron metabolism, influenced by cell line, tissue origin, and mechanism of ferroptosis induction (GPX4-independent versus dependent).

Many tumors including CRC, are characterized by elevated iron accumulation^14–17^, which suggests mechanisms to buffer excess redox-active iron to avoid cytotoxicity. CRC may uniquely have an adaptive mechanism to attenuate LIP size to limit oxidative damage which HCT116 and SW480 cells utilize. Identifying and targeting these iron-buffering pathways could represent an approach to increase the tumor sensitivity to ferroptosis-inducing therapies. Furthermore, the minimal transcriptional response of iron homeostasis genes (TRFC, DMT1, FTH1, FPN) suggests that ferroptosis initiation in these cells does not trigger broad changes in iron homeostasis, at least within the early time points examined. Taken together these results highlight an important distinction: while ferroptosis in CRC cells requires iron, the endogenous regulation of the LIP may not be the primary determinant of ferroptotic potentiation.

The compartmentalization of iron in subcellular compartments may also contribute to ferroptosis in ways not captured by bulk cytosolic LIP measurements. It has previously been shown that erastin-induced ferroptosis increases the LIP in the endoplasmic reticulum and lysosome in HT1080 cells^43^. Additionally recent work by Cañeque et al. demonstrates that activating lysosomal iron triggers ferroptosis in cancer cells, particularly affecting drug-tolerant persister populations^44^. This finding suggests that compartment-specific iron pools, such as lysosomal iron, play a pivotal role in ferroptosis regulation. Our observations are consistent with the idea that the cytosolic LIP remains largely stable during ferroptosis and could suggest that iron compartmentalization or redistribution, rather than changes in total cytosolic iron levels, plays a critical role in ferroptosis sensitivity. Further studies are needed to determine whether iron compartmentalization within subcellular compartments plays a role in CRC.

In summary, our study reveals that while exogenous iron can synergize with ferroptosis inducers in CRC, the LIP does not increase endogenously during ferroptosis. These findings redefine the assumed role of LIP dynamics in ferroptosis potentiation. Future studies should investigate whether targeting iron compartmentalization or buffering systems can enhance ferroptosis sensitivity in CRC and other iron-dependent cancers.

## Methods

### Drugs

RSL3, JKE1674, and IKE were prepared fresh from powder prior to use and diluted in DMSO. RSL3, JKE1674, and IKE were stored with a stock concentration of 10 mM in DMSO at -20°C. TRX-PURO was stored at a stock concentration of 4 mM in DMSO at -20°C. All final concentrations for experiments were prepared in cell culture media.

### Cell lines

Human intestinal colon cancer cell lines HCT116, SW480, RKO along with human embryonic kidney cells HEK293T, and fibrosarcoma cells HT1080 were utilized. All cell lines were cultured in complete DMEM medium supplemented with 10% fetal bovine serum and 1% antibiotic/antimycotic solution, maintained at 37°C in an atmosphere of 5% CO_2_ and 21% O_2_.

### Cell growth and synergy assays

Cells were plated between 250-500 cells/well and were allowed to adhere overnight, imaged for a day 0 reading and then immediately treated as indicated in the figure legend. Images were acquired 72 hours after treatment for final reading. Imaging was done on the Cytation 5 Imaging Multi-Mode reader with attached BioSpa from Agilent BioTek. Cytation software was used to quantify the cell count.

Analysis was performed by normalization to cell number at first reading (day 0) - all wells plated were averaged for analysis. Graphs were plotted using Prism with error bars representing mean +/- standard deviation.

To determine synergy between two different compounds, technical replicates of control and drug treatment wells within a plate were averaged. Fold changes relative to the averaged vehicle treatment group were calculated and then converted to a percentage to represent percent viability. The percent viability of the control groups (Vehicle, 250 μM, 500 μM, 1000 μM FAC) were then averaged to use as control values in the synergy analysis. These values were then used to determine synergy by the Bliss independence dose-response surface model using the SynergyFinder+ web application^14^.

### qPCR

Cell lines were treated for 16 hours and RNA was extracted using the Trizol reagent. RNA yield was then quantified using a Nanodrop. 1 μg of RNA was reverse transcribed to cDNA using the Invitrogen SuperScriptTM III First-Strand Synthesis System. After cDNA was collected Real time (RT) PCR reactions were done using three technical replicates for each sample. Then cDNA gene specific primers and SYBR green master mix were combined and then run on the Applied BioSystems QuantStudio 5 Real-Time PCR System. GAPDH was used as the housekeeping gene to calculate fold-change of the genes using the ΔΔCt method.

DMT1 F: CCTGTGGCTAATGGTGGAGTTGG

DMT1 R: GGAGATTGATGGCGATGGCTGAC

TRFC1 F: AGTTGAACAAAGTGGCACGAG

TRFC1 R: GCAGTTGGCTGTTGTACCTC

FTH1 F: TCCTACGTTTACCTGTCCATG

FTH1 R: GTTTGTGCAGTTCCAGTAGTG

FPN F: CACAACCGCCAGAGAGGATG

FPN R: CACATCCGATCTCCCCAAGT

### Western Blot

Cells were seeded in a 12-well plate each condition and allowed to adhere overnight. Cells were plated to reach ∼0.4×10^6 cells/well at time of harvest. After adherence cells were treated for 16 hours with RSL3, JKE1674, IKE, and with or without TRX PURO (1 mM). Cells were then lysed with RIPA assay buffer with added protease (1:100 dilution; MilliporeSigma) and phosphatase (1:100 dilution; Thermo Fisher Scientific) inhibitors. Lysates were quantified by BCA protein assay kit (Pierce – Thermo Fisher Scientific) and normalized for loading. Solubilized proteins were resolved on 10% SDS-polyacrylamide gels and transferred to nitrocellulose membrane, blocked with 5% milk in TBST, and immunoblotted with the indicated primary antibodies: Actin (1:1000) (Proteintech 66009-1-Ig), and α-puromycin (1:1000) (PMY-2A4). Horseradish peroxidase–conjugated secondary antibodies anti mouse (1:2500) were purchased from Cell Signaling (7076, 7074) and ThermoFisher Scientific (A15999). Immunoblots were imaged using the iBright imaging system.

### Flow Cytometry

After treatment with TRX-PURO (1 μM), RSL3 (0.25, 0.5, 1 μM), JKE (0.625, 1.25, 2.5 μM), or IKE (0.625, 1.25, 2.5 μM) for 16 hours cells were harvested using Trypsin and quenched with FACS buffer (PBS + 3% BSA + 1 u. They were then pelleted, washed, and resuspended in PBS in a 96 well round-bottom plate.<u>C</u>ells were then stained with live fixable blue (1:2500; diluted in PBS) (Invitrogen L23105) for 10 minutes, quenched with FACS buffer, and centrifuged at 600xg for five minutes. Cell pellets were washed once with PBS and then fixed with 4% paraformaldehyde on ice for 30 minutes. After fixation, the cells were stained overnight with Alexa647 anti-puromycin conjugated antibody (1:500 dilution) (BioLegend 381508) in permeabilization buffer (Invitrogen 00-8333-56) while protected from light.

### Statistical Analysis

Results are expressed as the mean +/- standard error of the mean for all figures unless otherwise noted. Significance between 2 groups was tested using a 2 tailed unpaired t test. Significance among multiple groups was tested using a one-way ANOVA. For all ANOVAs the Tukey-Kramer test was used for posthoc multiple comparison. GraphPad Prism 10.0 was used for the statistical analysis. Biorender was used to create all the schematics. Statistical significance is described in the figure legends as: ∗p < 0.05, ∗∗ p < 0.01, ∗∗∗ p < 0.001, ∗∗∗∗ p < 0.0001.

## Author Contributions

VP, PPH, YMS: conceived and designed the study; VP, DR, SS, PPH YMS: developed the methodologies; VP, DR, ZHL, NKD, LZ, KB: acquired the data; VP, DR, PPH, YMS: analyzed and interpreted the data; ARR provided TRX-PURO; VP, PPH, YMS: supervised the study and wrote the manuscript; and all authors: read and approved the final manuscript.

## Acknowledgements and Funding

The authors would like to acknowledge the members of the Hsu and Shah laboratories for project feedback and technical expertise. This work was supported by NIH grants R01CA148828, R01CA245546, and R01DK095201 to YMS. The work was supported by the University of Michigan Comprehensive Cancer Center Core grant P30CA046592. VP is also supported by NCI diversity supplement as part of the parent R01CA245546.

## Supplemental Figure Legends

**Supplemental Figure 1:**
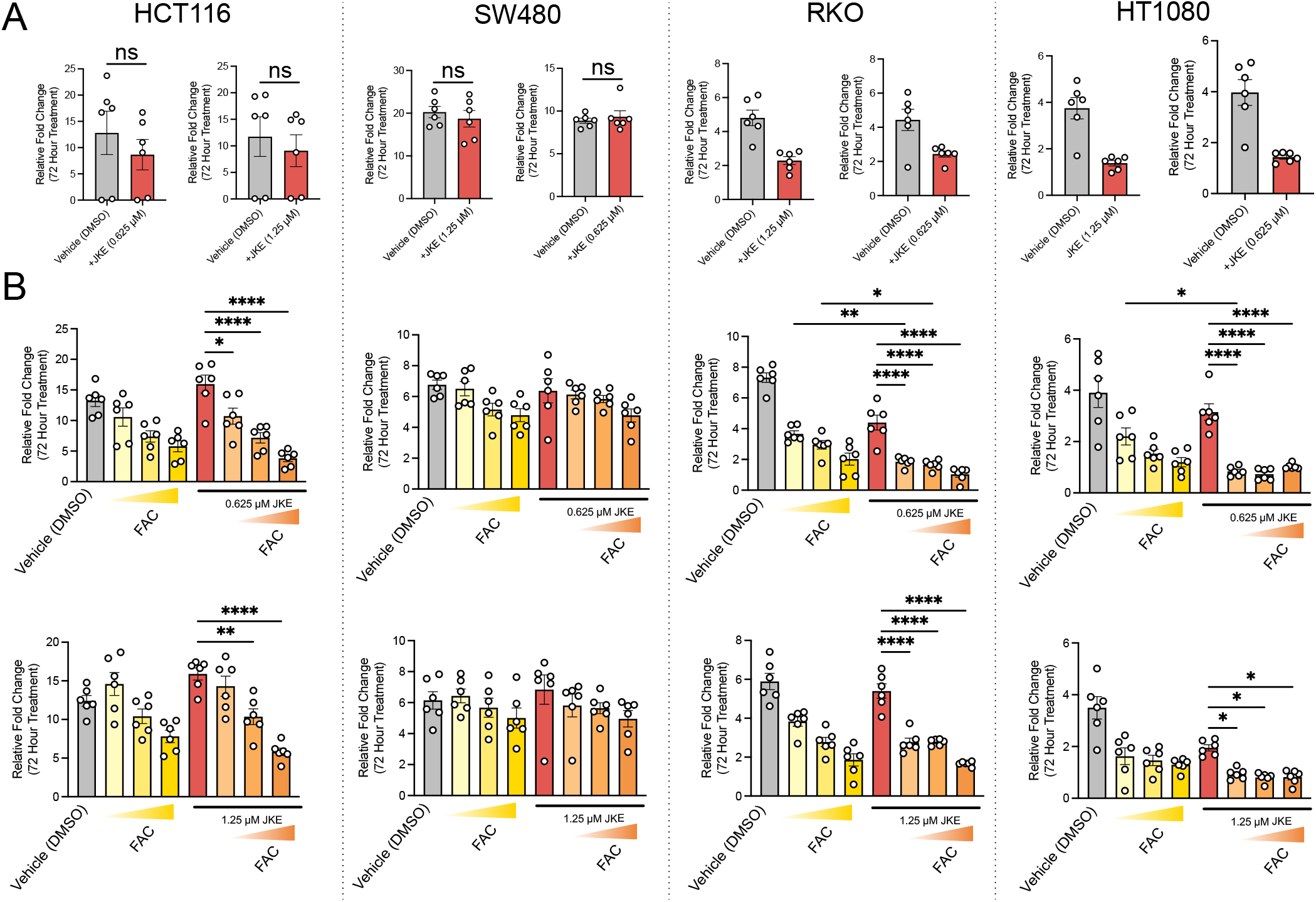
Exogenous iron sensitizes CRC cells to ferroptosis at lower concentrations of JKE. (A) Cell viability of HCT116, SW480, RKO, and HT1080 cells following treatment with 1.25 and 0.625 μM JKE1674 after 72 hours Cell viability of HCT116, SW480, RKO, and HT1080 cells following treatment with 1.25 and 0.625 μM JKE1674 with the addition of 250, 500, 100 μM FAC after 72 hours ∗p < 0.05, ∗∗p < 0.01, ∗∗∗p < 0.001, ∗∗∗∗

**Supplementary Figure 2:**
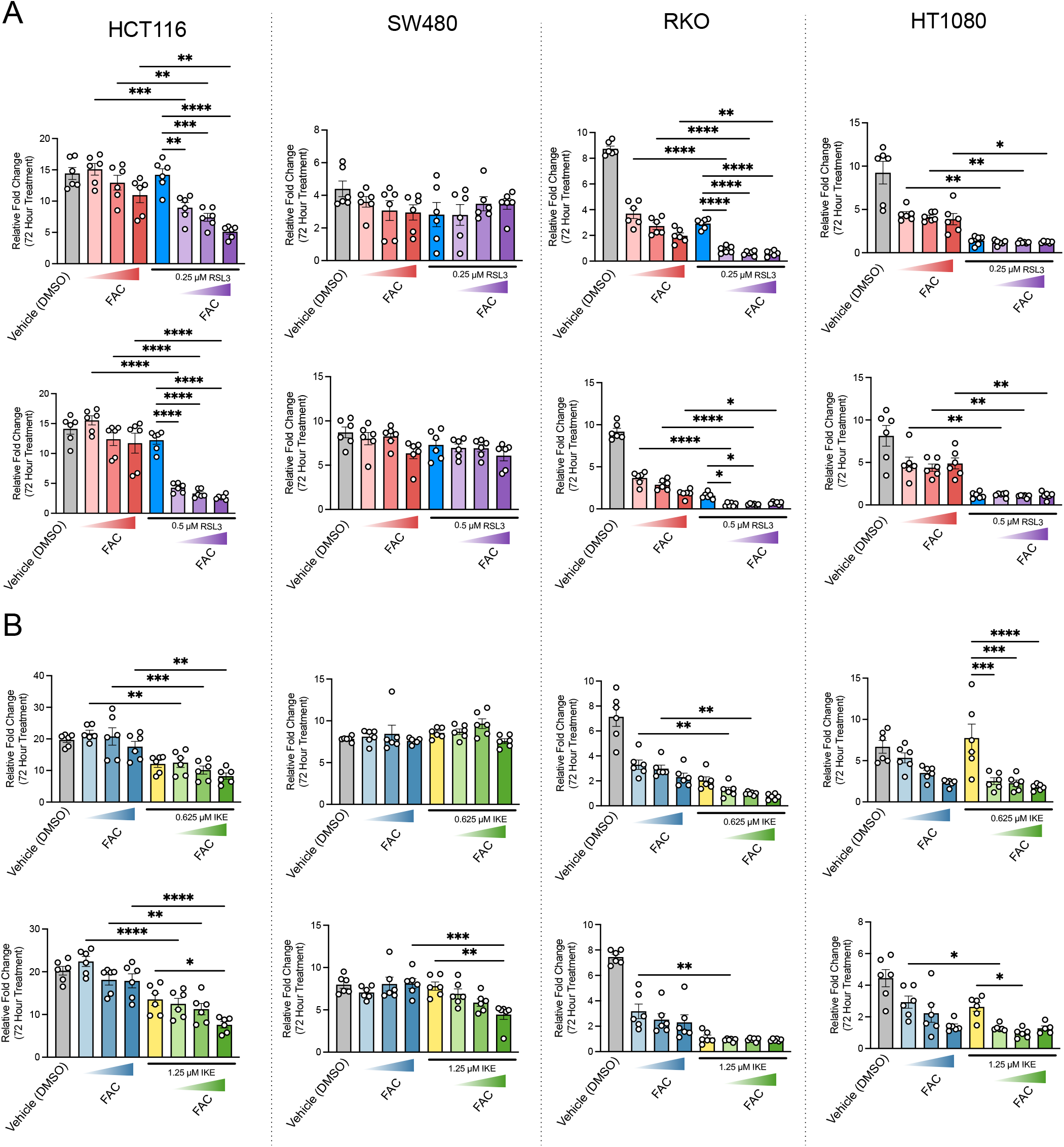
Exogenous iron sensitizes CRC cells to ferroptosis at lower concentrations of RSL3 and IKE. (A) Cell viability of HCT116, SW480, RKO, and HT1080 cells following treatment with 1.25 and 0.625 μM RSL3 after 72 hours (B) Cell viability of HCT116, SW480, RKO, and HT1080 cells following treatment with 1.25 and 0.625 μM IKE with the addition of 250, 500, 100 μM FAC after 72 hours ∗p < 0.05, ∗∗p < 0.01, ∗∗∗p < 0.001, ∗∗∗∗p < 0.0001 unless otherwise indicated. Data are presented as mean ± SEM.

**Supplementary Figure 3:**
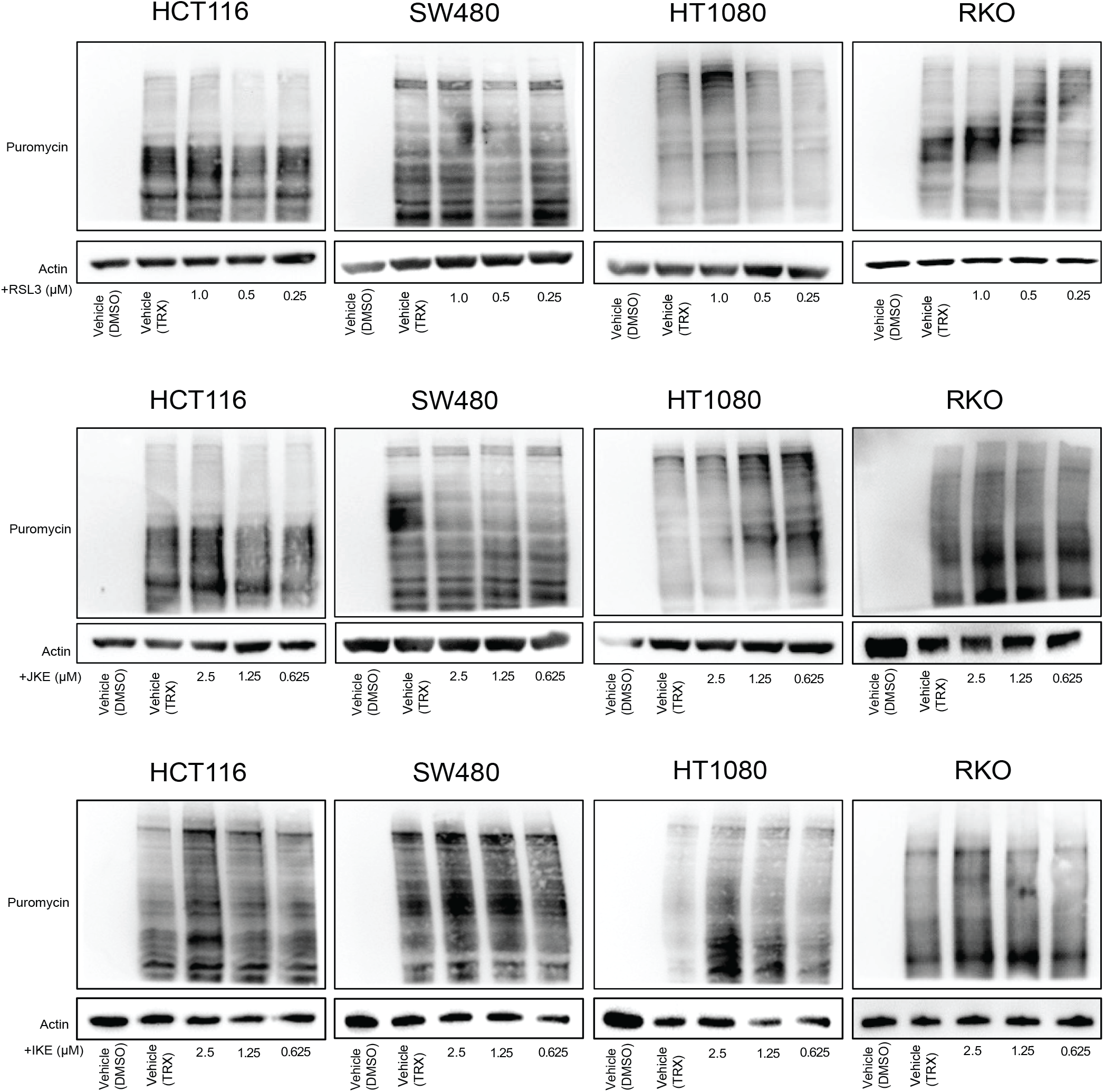
Measuring changes in cytosolic labile iron using TRX-PURO in response to ferroptotic inducers by western blotting. (A) Western blot of HCT116, SW480, HT1080, RKOs to assess labile iron pool concentration after treatment of 1, 0.5, 0.25 μM RSL3 and TRX-PURO (1 μM) (B) Western blot of HCT116, SW480, HT1080, RKOs to assess labile iron pool concentration after treatment of 2.5, 1.25, 0.625 μM JKE and TRX-PURO (1 μM) (C) Western blot of HCT116, SW480, HT1080, RKOs to assess labile iron pool concentration after treatment of 2.5, 1.25, 0.625 μM IKE and TRX-PURO (1 μM)

## Notes

### Competing Interest Statement

The authors have declared no competing interest.

### Summary of Updates

Figure 4 revised; added one more supplemental figure

